# Developing synthetic auxin-based trans-kingdom signaling

**DOI:** 10.64898/2026.11.03.764009

**Authors:** Madison Furnish, Emmanuel J Kokarakis, Dahlia Kopycienski, Bjorn Thune-Larsen, Daniel C Ducat, Arjun Khakhar

## Abstract

Synthetic trans-kingdom signaling systems with tunable transduction properties provide a unique avenue to explore how communication between organisms coordinate behaviors without the confounding complexity of natural systems. However, existing trans-kingdom signaling systems suffer from either toxic signal molecules, a need for rare precursors for signal generation, or transduction machinery which is either species-specific or cannot be easily tuned. This limits their capacity to be broadly applied across kingdoms and their utility as a model system to study trans-kingdom interactions. In this work we elucidate the design principles for a trans-kingdom signaling system based on the plant phytohormone, Indole 3-Acetic Acid (auxin), that overcomes these challenges. We demonstrate how auxin biosynthesis can be engineered into a model sender organism, the cyanobacteria *Synechococcus elongatus* PCC 7942, and how this signal can be tuned with low leak and high dynamic range. We also show these signals can be connected to gene expression using tunable synthetic auxin-responsive signaling systems across multiple kingdoms. We demonstrate that *S. elongatus* generated auxin signals can tune gene expression in co-cultured *Saccharomyces cerevisiae* via auxin-induced degradation. Additionally, we show that when the *S. elongatus* are associated with the roots of the model plant, *Arabidopsis thaliana*, the auxin they generate can control gene expression via auxin-responsive synthetic transcription factors expressed from the plant’s genome. Taken together, our results demonstrate the design principles for a tunable auxin-based trans-kingdom signaling system that can be broadly applied across species.

## Introduction

Chemical signals are often used to carry information between organisms to coordinate behaviors. In some situations, this trans-kingdom signaling facilitates mutualisms, as are observed in lichen^1,2^ or gut microbiomes^3^, and in others it forms the basis of pathogenesis, as can be seen in infections of plants by *Agrobacterium*^4^ or rodents by *Toxoplasma gondii*^5^. Thus, understanding how quantitative differences in trans-kingdom signaling shape organismal interactions is critical to uncovering the mechanistic basis of the interactions that underpin ecosystems. Achieving this understanding requires the capacity to tune aspects of signaling such as signal strength, perception sensitivity, or downstream targets of regulation, and then characterize changes in interactions. This is challenging to do via studies of complex natural variation where multiple aspects of signaling change in tandem. The development of synthetic trans-kingdom signaling systems with tunable signaling parameters as a model system to study these questions is a promising avenue to overcome this challenge^1^.

An ideal synthetic trans-kingdom signaling system would include a sending module that could be engineered to enable signal generation with a tunable strength, and a receiver module in which the transduction properties and downstream regulation can be flexibly programmed. Additionally, these modules would be composed of components that are functional across species, and the signal would be minimally toxic, to enable broad application across different species pairs. While prior work has elucidated several possible signaling systems that could be used to communicate across kingdoms^6^, they generally suffer from either toxic signal molecules, a need for precursors from unusual secondary metabolism to generate the signal in the sender species, or transduction machinery which is either species-specific or cannot be easily tuned.

A trans-kingdom signaling system based on the phytohormone Indole 3-Acetic Acid (IAA or auxin) has the potential to overcome these issues. The central role that auxin plays in plant signaling^7^ makes it ubiquitous in the environment and hence it is rarely toxic to organisms at physiological concentrations. Prior work has elucidated a two enzyme pathway from *Agrobacterium tumefaciens* that uses the ubiquitous amino acid tryptophan as a substrate to biosynthesize auxin^4^. This pathway consists of the enzyme *iaaM*, which converts tryptophan into the auxin precursor, indole-3-acetamide (IAM), and the enzyme *iaaH*, which converts IAM into auxin. It has been functionally expressed in a range of organisms, including bacteria, fungi, and plants^8,9^ which demonstrates its broad utility as signal generation module for a trans-kingdom signaling system.

Complementary studies have shown that synthetic signaling systems which respond to auxin can be constructed using plant derived auxin-triggered degradation domains, called auxin degrons^8,10^. These systems operate through the fusion of degrons either directly to proteins of interest^8^ or to synthetic transcription factors^11^ that can be flexibly programmed to regulate the expression of genes of interest. In the presence of auxin and an auxin-responsive F-Box protein, such as AFB2, the protein fused to the auxin degron is polyubiquitinated by the E3 ubiquitin ligase machinery and subsequently degraded in the proteosome. This enables auxin signals to be coupled to gene expression, either directly via degradation in the case of degron fusion, or indirectly via degradation of an auxin-responsive synthetic transcription factor which relieves regulation. These auxin-degron based synthetic signaling systems have been demonstrated to have tunable auxin responsivities, programable regulation of downstream targets, and functionality across diverse organisms such as fungi and plants^8,11^. These properties make them an ideal receiver module for a trans-kingdom signaling system.

In this work we implement an auxin signal generation module in the cyanobacterium *Synechococcus elongatus* PCC 7942 and characterize its auxin generation over time, its tunability, and the constraints on its function. We then explore whether the auxin signals it generates can be coupled to synthetic signaling systems expressed in organisms from other kingdoms, including a fungus, *Saccharomyces cerevisiae* (yeast), and a plant, *Arabidopsis thaliana*. Through this work we hope to elucidate the design principles for a tunable auxin-based synthetic trans-kingdom signaling system that can be broadly applied across species for future, bottom-up studies on the role of signaling in shaping organismal interactions.

## Results

### Expression of *iaaH* and *iaaM* in *S. elongatus* enables auxin biosynthesis

To enable trans-kingdom signaling, we expressed the auxin biosynthesis pathway from *A. tumifaciens* in *S. elongatus.* We created a plasmid encoding the *iaaH* and *iaaM* genes controlled under a previously validated theophylline responsive riboswitch **(Fig 1A)** and integrated this expression cassette into the *S. elongatus* genome at a defined neutral site locus. Validated transformants were cultured in BG11 media, induced with 0.5mM theophylline, and auxin secretion was evaluated over time via LC-MS relative to mock controls. We observed a steady increase in auxin concentration in the media over time in the theophylline treated cultures **(Fig 1BC)**, whereas auxin did not accumulate in uninduced cultures, indicating that *S. elongatus* does not have native pathways for auxin biosynthesis. We tracked cell growth of these cultures over time, observing that theophylline-treated cultures exhibited slightly lower cell density at later time points, however these differences were not significant **(Fig 1B)**. We hypothesize that utilization of endogenous tryptophan by *iaaH*, the metabolic load of enzyme expression, or the theophylline inducer itself could marginally impact growth rates. By normalizing the measured auxin concentrations to cell density, we obtain an estimate of specific productivity of auxin, which peaks at 24 hours post induction **(Fig 1D)**. Together these results demonstrate that expression of *iaaH* and *iaaM* in *S. elongatus* can enable auxin biosynthesis and that this biosynthesis increases for 24 hours post induction and then attenuates over time.

**Fig 1.**
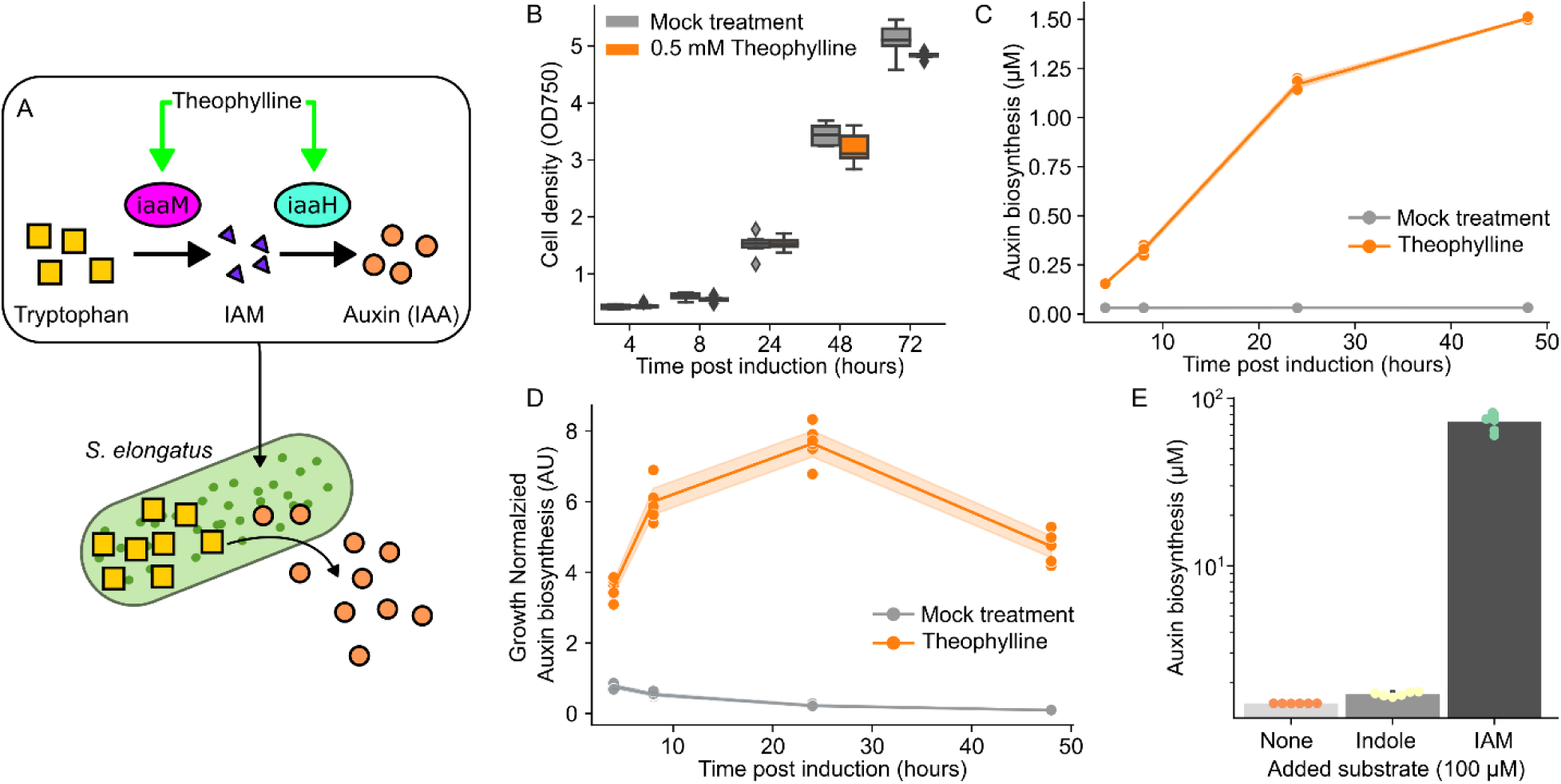
A) Schematic depicting the two-enzyme pathway activated via theophylline induction which converts tryptophan (yellow squares) into auxin (orange circles) in *S. elongatus.* B) Box plots summarizing cell growth over time (n=6), measured as OD_750_, for an *S. elongatus* strain expressing the circuit shown in (A) treated with either 0.5 mM theophylline (orange) or a mock treatment (gray). C) Plot summarizing auxin concentration in the culture supernatant over time from a culture of the strain described in (A) after induction with either 0.5 mM theophylline (orange) or a mock treatment (gray). Shaded area around lines connecting means represent the 95% confidence interval (n=6). D) Plot summarizing auxin concentration normalized by growth for the same cultures shown in (C). Shaded area around lines connecting means represent the 95% confidence interval (n=6). E) Bar plots summarizing auxin concentration in the supernatant of cultures (n=6) of the strain described in (A) treated with 0.5 mM theophylline and either 100 µM IAM (teal dots), 100 µM Indole (yellow dots), or a mock (orange dots). Error bars represent standard deviation.

### The generation of IAM is the rate limiting step in auxin signal generation

While our results thus far show that auxin biosynthesis is possible in *S. elongatus,* the amount of auxin generated was lower than the sensitivity threshold for most auxin responsive synthetic signaling systems^8^. We hypothesized that this was due to either due to a limited pool of tryptophan or poor flux through a key enzymatic step. To understand where the rate limiting step in auxin biosynthesis was, we grew the engineered strain in BG11 media containing 0.5 mM theophylline to induce the pathway and either 100 µM Indole, 100 µM IAM, or a mock treatment. Tryptophan is not membrane permeable and so cannot be directly taken up from the media by *S. elongatus*. However, indole is a membrane-diffusible precursor to tryptophan biosynthesis, providing an alternative to increase the bioavailable pool of the substrate. We then grew these strains for 48 hours and measured auxin in the supernatant as described previously. We observed no significant effect from addition of indole, but a multiple order-of-magnitude increase in auxin biosynthesis upon addition of IAM to the media **(Fig 1E)**. By adding IAM to the media, an auxin signal can be generated that is well above the detection threshold of synthetic auxin signaling systems. The lack of impact of adding indole could be explained by either a low conversion efficiency of indole into tryptophan by *S. elongatus*, feedback regulation which prevents a large bioavailable pool of tryptophan in the cell^12^, or a low conversion efficiency of tryptophan to IAM by *iaaM*. Together, these results indicate that the conversion of tryptophan to IAM is the most likely rate-limiting step in the production of auxin by our strain, and that media supplementation by IAM enables strong auxin signal generation.

### Expression of *iaaH* in *S. elongatus* grown in IAM supplemented media enables tunable auxin signal generation

Given the mild penalty from expression of the auxin biosynthesis pathway on *S. elongatus* growth, and the likelihood that IAM availability was limiting auxin production, we reasoned that decoupling auxin signal generation from IAM biosynthesis would be useful. To achieve this, we generated alternative strains of *S. elongatus* solely for inducible expression of *iaaH* using a similar strategy as described for the full biosynthesis pathway described above (**Fig 2A**). This strain was then grown in BG11 media supplemented with 100µM IAM and a range of theophylline concentrations spanning 0 to 1 mM, while measuring both cell growth and chlorophyll *a* content as a proxy for maintenance of normal photosynthetic capacity. We observed no growth penalties for theophylline inducer concentrations up to 0.5 mM with either condition (**Fig 2B, Supp Fig 1,2**). A slight growth penalty was observed at 1 mM theophylline, which is consistent with documented toxicity of theophylline at higher concentrations^13–15^. Total auxin bioproduction and cell density normalized production rate were correlated with the level of theophylline inducer at concentrations up to 1 mM **(Fig 2D, Supp Fig 3)**, though theophylline concentrations above 0.05 mM showed diminishing returns of auxin production, suggesting that the maximal effective expression of *iaaH* was achieved at inducer concentrations well below the toxicity of 1mM theophylline. Additionally, total auxin measured in the culture supernatant peaked at 48 hours for higher induction levels of *iaaH* (>0.05 mM theophylline), while it continued to accumulate up to 72 hours in cultures induced with less than 0.005 mM theophylline **(Fig 2C)**. Consistently, when normalized to cell density, the rate of auxin bioproduction peaked at 24 hours under higher *iaaH* induction (≥0.05 mM theophylline) and peaked at 48 hours at lower induction (≤0.005 mM theophylline; **Supp Fig 4**). LC-MS analysis of supernatant from later timepoints indicates that declining auxin productivity over time is due to the depletion of the exogenously added IAM substrate (**Fig 2E**). Together, these results demonstrate that an *S. elongatus* with inducible expression of *iaaH* which is grown in IAM supplemented media can generate a strong and tunable auxin signal.

**Fig 2.**
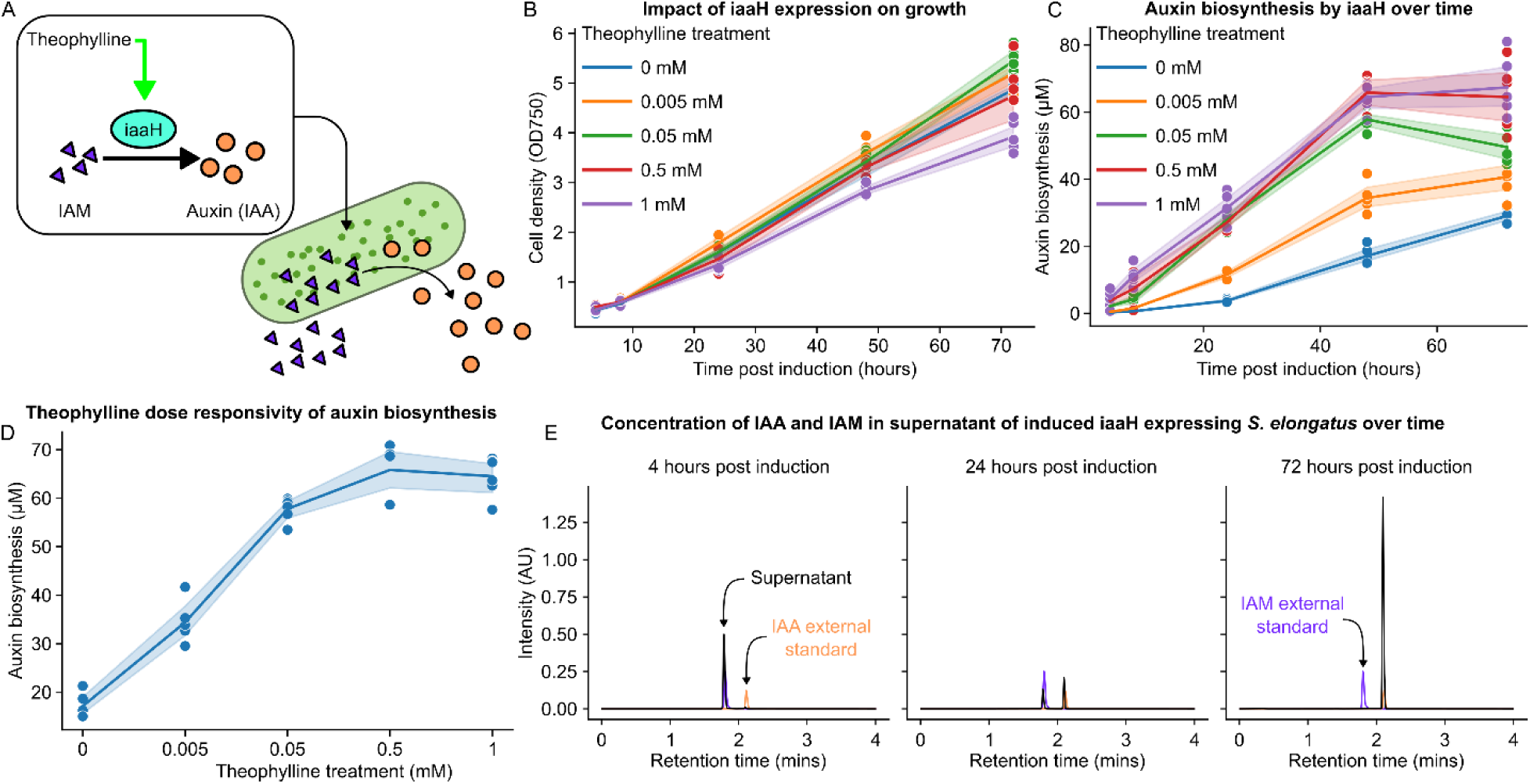
A) Schematic depicting theophylline induced expression of *iaaH* in *S. elongatus* which leads to conversion of IAM in the media (purple triangles) to auxin (orange circles). B) Plot summarizing cell growth over time, measured as OD750, of the strain described in (A) in media containing 100 µM IAM post induction with varying concentrations of theophylline. Shaded area around lines connecting means represent the 95% confidence interval (n=6). C) Plot summarizing auxin concentration over time in the supernatant of cultures of the strain described in (A) in media containing 100 µM IAM and varying concentrations of theophylline. Shaded area around lines connecting means represent the 95% confidence interval (n=6). D) Plot summarizing the auxin concentration at 48 hours post induction in the supernatant of cultures of the strain shown in (A) with varying levels of theophylline induction. Shaded area around lines connecting means represent the 95% confidence interval (n=6). E) Chromatograms in black summarizing LC-MS characterization of the supernatant from cultures of the *S. elongatus* strain shown in (A) induced with 100 µM IAM and 0.5 mM theophylline after 4, 24, and 72 hours. The chromatograms of an IAA (orange) and IAM (purple) external standard are overlaid on each plot.

### Tuning signal strength using IAM rather than theophylline enables lower leak and broader dose response

In the preceding auxin biosynthesis experiments under supplemental IAM addition, we observed significant concentrations of auxin produced even with no theophylline inducer **(Fig 3A)**, suggesting leaky expression of *iaaH* from the riboswitch element. Indeed, the final auxin bioproduction at 72 hours was only 2.32-fold higher when maximally induced (1mM theophylline) versus uninduced control (29.1 µM compared to 67.4 µM; **Fig 3A**). This led us to explore whether the dynamic range of the system could be increased by holding *iaaH* expression constant (with 1 mM theophylline) while varying the addition of IAM substrate. Indeed, by varying IAM supplementation in *S. elongatus* cultures from 0 µM to 100 µM we observe auxin concentrations from 0.021 µM to 67.4 µM respectively (**Fig 3B**), representing a 3089-fold change between the uninduced and induced condition. Similarly, holding theophylline at the non-toxic concentration of 0.5 mM, we observe a dose dependent increase in auxin generated with increasing concentrations of IAM over a dynamic range spanning three orders of magnitude (**Fig 3C**). These results demonstrate that titrating IAM concentration is a superior method to tune the strength of the auxin signal generated by *S. elongatus* for trans-kingdom signaling.

**Fig 3.**
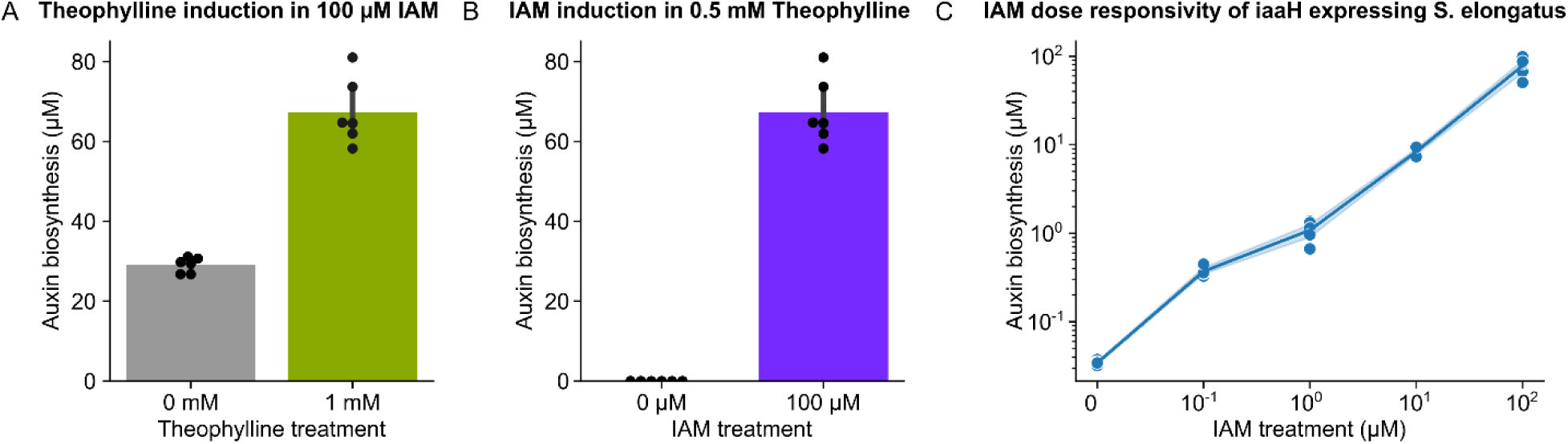
A) Bar plot summarizing auxin concentration at 72 hours post induction in the supernatant of cultures of *S. elongatus* expressing theophylline induced *iaaH* treated with 100 µM IAM and different concentrations of theophylline (n=6). B) Bar plot summarizing auxin concentration at 72 hours post induction in the supernatant of cultures of *S. elongatus* expressing theophylline induced *iaaH* treated with 0.5 mM theophylline and different concentrations of IAM (n=6). C) Plot summarizing auxin concentration at 72 hours post induction in the supernatant of cultures of *S. elongatus* expressing theophylline induced *iaaH* treated with 0.5 mM theophylline and a range of IAM concentrations. Shaded area around lines connecting means represent the 95% confidence interval (n=6).

### Auxin-based trans-kingdom signaling enables communication between cyanobacteria and yeast in co-culture

Our results so far establish that *S. elongatus* expressing a theophylline induced *iaaH* gene can produce a tunable auxin signal via titration of IAM in the media with minimal growth defects. Next, we wanted to validate the feasibility of using this system for trans-kingdom signaling. Prior studies have shown that *S. elongatus* and *Saccharomyces cerevisiae* (yeast) are compatible with growth in a co-culture^16–18^. We decided to build on this work by exploring if the auxin signal generated by *S. elongatus* could be coupled to gene expression in yeast. To do this we used a strain of *S. cerevisiae* expressing a previously validated synthetic auxin signal transduction system regulating a reporter whose output can be ratiometrically normalized^19^. In this system an auxin degron from the *Arabidopsis thaliana INDOLE ACETIC ACID-INDUCED PROTEIN 17* (IAA17) protein in plants is fused to a Venus fluorescent reporter (YFP). In the presence of auxin this protein associates with a plant F-Box protein, AFB2, fused to a mScarlet fluorescent reporter (RFP) that is co-expressed from the same polycistronic messenger RNA (**Fig 4A**). This association triggers auxin-induced polyubiquitination and subsequent proteasomal degradation of the IAA17-YFP reporter. Cell-to-cell variation in the expression of YFP, which would normally confound auxin responsive changes in YFP signal, can be normalized using the constitutively expressed RFP.

**Fig 4.**
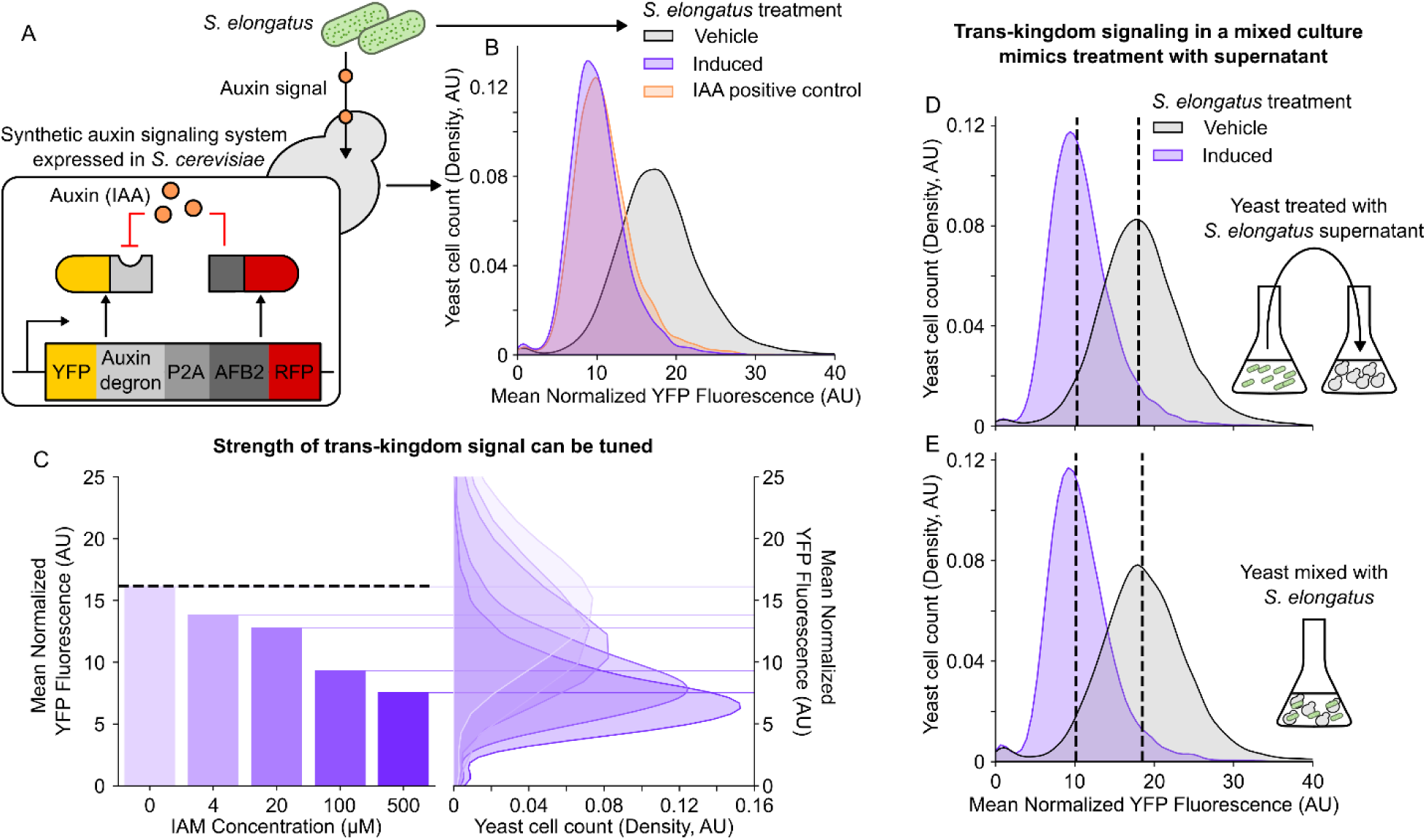
A) Schematic describing how an auxin signal generated by engineered *S. elongatus* is sensed by a synthetic signaling system in *S. cerevisiae* consisting of a YFP fused to an auxin degron and an RFP fused to an auxin receptor, AFB2, both of which are expressed from a polycistronic mRNA. B) Kernel density estimate plots (smoothed histograms) summarizing flow cytometry-based characterization of mean YFP fluorescence normalized by RFP fluorescence of approximately 10,000 yeast cells described in (A) treated with supernatant from cultures of *S. elongatus* with inducible auxin biosynthesis. Plot colors correspond to *S. elongatus* treatment conditions. C) Bar plots summarizing mean normalized YFP fluorescence of approximately 10,000 yeast cells described in (A) treated with supernatant from cultures of *S. elongatus* with inducible auxin biosynthesis that were incubated with the indicated concentrations of IAM. Paired Kernel density estimate plots (right) depict the distribution of fluorescence of the population around the mean. Dashed black line depicts the mean normalized YFP fluorescence of yeast treated with supernatant from *S. elongatus* grown without theophylline or IAM. D,E) Kernel density estimate plots summarizing flow cytometry-based characterization of mean normalized YFP fluorescence of approximately 10,000 yeast cells described in (A) are shown following either (D) treatment with supernatant from *S. elongatus* incubated with 0.5 mM theophylline and 100 µM IAM or (E) direct mixing of yeast cells of with the same *S. elongatus* strain incubated in the same treatment. Dashed black lines represent the median of the populations.

To isolate the impact of the auxin signal from other co-culture dependent effects on gene expression in the yeast strain, we initially tested the response of the yeast to supernatant from the *S. elongatus* culture. We cultured the *S. elongatus* in BG11 media with 1mM of theophylline and either 100 µM IAM or DMSO as a negative control for 72 hours then added the filtered supernatant to a culture of the yeast strain in log phase growth. We also set up a positive control, in which we added a saturating concentration (100 µM) of IAA to a log phase yeast culture. The yeast was then allowed to grow for another 3 hours under induction to achieve a steady state after which single cell fluorescence measurements were made using a flow cytometer. We observed a 37.1% decrease in the mean normalized YFP signal of the yeast treated with supernatant from *S. elongatus* grown in theophylline and IAM compared to yeast treated with supernatant from the negative control (**Fig 4B**). This was approximately the same as the 34.7% decrease in the mean normalized YFP signal observed in the positive control (**Fig 4B**), demonstrating that the auxin signal generated by the *S. elongatus* could maximally induce auxin-responsive protein degradation in yeast.

Next, we explored whether sub-saturating auxin signals generated by *S. elongatus* can tune gene expression in yeast in a dose-dependent manner. We repeated the experiment described above using supernatant from *S. elongatus* cultures treated with 1mM theophylline and IAM concentrations ranging between 0 µM and 500 µM. We observed a dose-dependent decrease in mean normalized YFP signal in response to increasing IAM concentrations the *S. elongatus* was treated with (**Fig 4C**). We observed a broad dynamic range in dose-responsivity to the auxin signal, spanning a fold change in the YFP signal of 8.4-fold between the 0 µM and 500 µM IAM treatments. We also observed minimal leaky degradation of YFP, as evidenced by a 0.18-fold change in YFP signal between the untreated yeast and those treated with 0 µM IAM *S. elongatus* culture supernatant (**Fig 4C**). While these results demonstrate effective dose-dependent trans-kingdom signaling between *S. elongatus* and yeast via IAM based induction, we wanted to confirm that this strategy was superior to inducing with theophylline, as indicated by the LC-MS based characterization of *S. elongatus*. We repeated the dose response experiment using supernatant from *S. elongatus* treated with 100 µM IAM and 5, 50, 100, and 1000 µM of theophylline. While we observed the expected decrease in mean normalized YFP signal at the higher theophylline concentrations, the dynamic range was 192% lower and the leak was 23.5% higher than when IAM was used for induction (**Supp Fig 5**).

Finally, we wanted to explore whether the auxin signals generated by *S. elongatus* can efficiently regulate gene expression in yeast when they are in the proximity of a co-culture. We grew the *S. elongatus* in media treated with 1mM theophylline and 100 µM IAM or DMSO as a negative control and then mixed it with a culture of the yeast in log phase. In tandem, we also mixed another culture of yeast with an equivalent amount of filtered supernatant, as in prior experiments. After 3 hours of growth, the florescence signal in the yeast was then measured at a single cell level by performing flow cytometry on these cultures. In the co-cultured samples, the forward and side scatter were used to separate fluorescent measurements of yeast from *S. elongatus.* We observed a nearly identical decrease in mean normalized YFP signal in both co-culture and supernatant treated yeast compared to the negative control (**Fig 4 D,E**), demonstrating that this trans-kingdom signaling system can operate in a co-culture setting. Together, these results show that efficient auxin-based trans-kingdom signaling can be implemented between *S. elongatus* and *S. cerevisiae*.

### Auxin-based trans-kingdom signaling enables communication between cyanobacteria and plant roots

Our results thus far demonstrate that our system can enable auxin-based signaling between *S. elongatus* and yeast. We next sought to explore if this system was portable enough to be extended to a different sender-receiver pair. Prior work has shown that *S. elongatus* can associate efficiently with plant roots^20^. Additionally, plants natively use auxin as signaling molecules to coordinate development, so it was expected that the *S. elongatus* generated signal would alter plant gene expression. However, we were interested in whether this auxin signal could be programmatically coupled to the expression of any gene of interest, rather than the specific ones that auxin natively regulates in plants. To achieve this, we used a transgenic line of *Arabidopsis thaliana* expressing an auxin responsive Cas9-based synthetic transcription factor, consisting of deactivated Cas9 fused to a repression domain from the TOPLESS protein and an auxin degron, targeted to repress the expression of a reporter whose functionality we have previously validated^11^. Here, a co-expressed guide RNA targets the Cas9-based synthetic transcription factor to the promoter driving the expression of fusion protein consisting of a nuclear localized fluorescent protein (Venus) fused to firefly luciferase, repressing its expression. In the presence of auxin, the transcription factor undergoes ubiquitination followed by proteasomal degradation, relieving repression and increasing reporter expression (**Fig 5A**).

**Fig 5.**
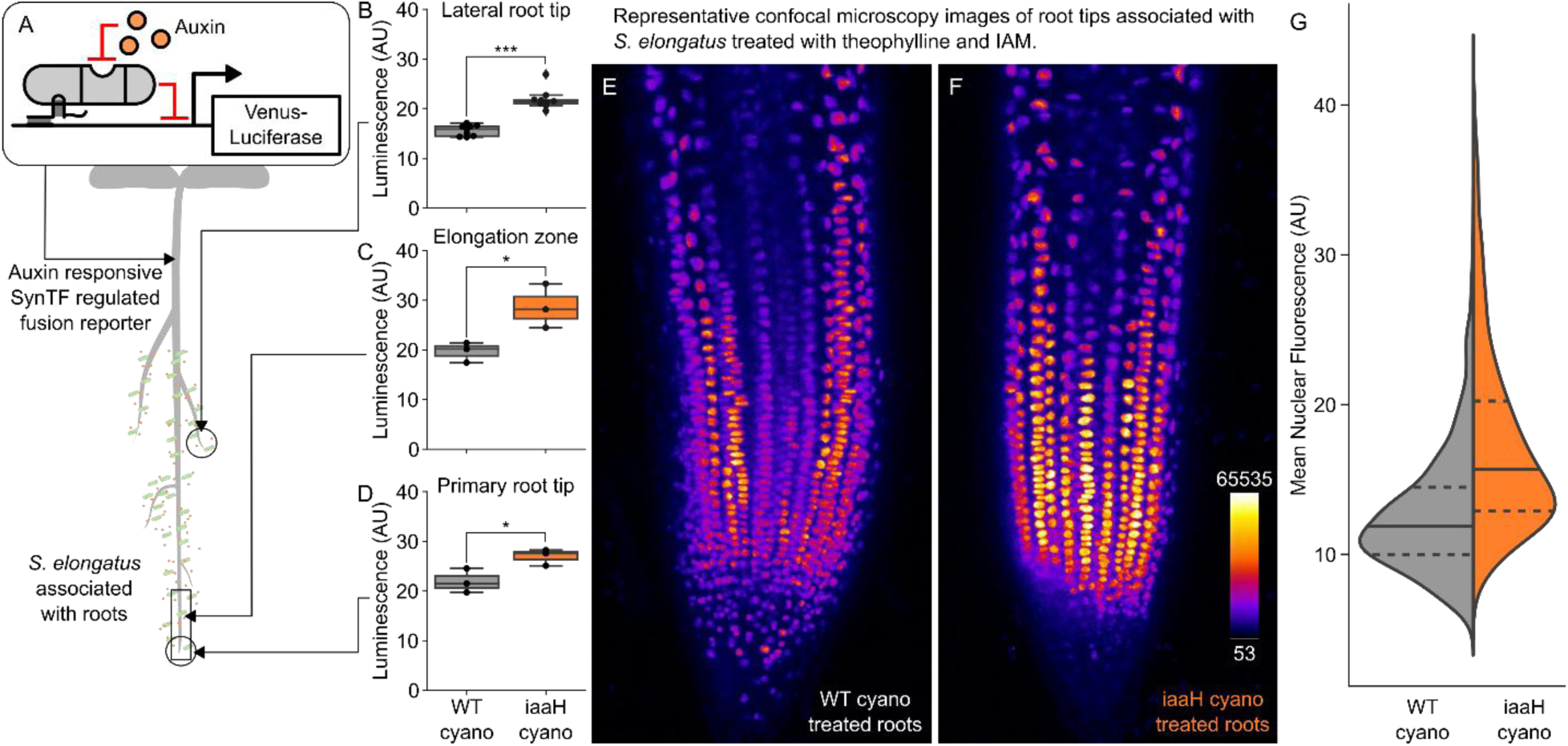
A) Schematic depicting auxin signals produced by root associated *S. elongatus* being transduced by an auxin responsive Cas9-based synthetic transcription factor to trigger expression of a nuclear localized fluorescent reporter fused to luciferase. B,C,D) Box plots summarizing bioluminescence signal in lateral root tips (B), elongation zones (C), or primary root tips (D) of plants expressing the circuit shown in A with either wildtype (gray) or *iaaH* expressing (orange) *S. elongatus* induced with 100 µM IAM and 0.5 mM theophylline associated with their roots. Every dot represents an independent biological replicate, * represents p<0.05, and *** represents p<0.0005. E,F) Representative maximum intensity projections of confocal microscopy images of the lateral root tips from plants expressing the circuit shown in A with either wildtype (E) or *iaaH* expressing (F) *S. elongatus* induced with 100 µM IAM and 0.5 mM theophylline associated with their roots. G) Stacked violin plot showing the distribution of mean fluorescence of nuclei from the microscopy shown in (E). Each histogram represents aggregated measurements from all segmented nucli in confocal images of three independent biological replicates, with the solid black line showing the mean, and the dashed black lines showing the quartiles of the population.

We explored whether the auxin signal generated by *S. elongatus* expressing *iaaH* was sufficient to modulate the expression of reporter expression in roots by growing *S. elongatus* in 100 µM IAM and 0.5mM theophylline as described above then inoculating the roots of seedlings of *A. thaliana* plants expressing the circuit described above grown vertically on 0.5xMS plates. A parallel set of control plants were treated with wildtype *S. elongatus* that were grown in the same conditions to control for the impact of IAM in the induction media. After 24 hours in long day growth conditions, the expression of the fusion reporter in treated roots was measured at the tissue level by treatment with 5µM D-luciferin followed by imaging-based quantification of bioluminescence. We observed significantly higher reporter signal in the elongation zone, primary root tip, and lateral root tips of seedlings treated with the auxin producing *S. elongatus* strain compared to roots treated with the wildtype control (**Fig 5B-D**).

We further characterized the auxin-triggered reporter de-repression at a cellular resolution in the root using confocal microscopy. Here lateral root tips from plants treated as described above were mounted on slides and imaged to measure the signal from the nuclear localized fluorescent reporter (**Fig 5E,F**). We then used a previously validated software package to segment individual nuclei and measure the mean fluorescence signal for each nucleus. These measurements across multiple root tips were then aggregated and compared to parallel measurements from plant roots treated with the wildtype control. We observed an increase in the mean nuclear fluorescence in plants treated with *iaaH* expressing *S. elongatus* compared to those treated with wildtype (**Fig 5G**). Together, these results demonstrate that auxin signals generated by *S. elongatus* can be transduced into changes in gene expression in plant roots, enabling trans-kingdom signaling.

## Discussion

Phytohormones like auxin play important roles in natural trans-kingdom signaling, both between microbes and plants as well as within microbial consortia. Creating synthetic versions of these signaling systems whose signal strength and transduction properties can be tuned will enable precision control of this important communication channel. In this work we elucidate the design rules for the implementation of a synthetic auxin-based trans-kingdom signaling system and demonstrate its utility for enabling communication across bacteria, plants, and fungi.

Our results show that the theophylline dependent expression of *iaaM* and *iaaH* is sufficient to enable inducible generation of auxin signals from the endogenous pool of tryptophan in *S. elongatus*. The universal availability of the amino acid substrate and the broad biocompatibility of auxin make this pathway an ideal signal generation module. However, we observed a relatively low concentration of auxin produced in initial engineered strains of *S. elongatus* and identified the generation of IAM from tryptophan to be the rate limiting step. We hypothesize that the low conversion of tryptophan is due to either poor catalysis of the *iaaM* enzyme used, feedback regulation limiting the pool of bioavailable tryptophan, or some combination of both. While outside the scope of this study, future work to increase auxin biosynthesis by identifying efficient IAM biosynthesis enzymes or increasing tryptophan availability through metabolic engineering would improve the dynamic range of this sending module.

We overcame poor auxin production by supplementing the media with IAM and engineering the *S. elongatus* to only express *iaaH* in response to the theophylline inducer. This enabled the tunable generation of a strong auxin signal with a large dynamic range. Interestingly, our characterization of the *iaaH* expressing strain revealed that the dynamic range could be substantially increased, via a reduction of leak in the off state, by tuning the auxin biosynthesis through modulation of IAM in the media rather than the theophylline inducer. The superiority of IAM over theophylline for tuning the strength of the auxin signal generated was further confirmed from our experiments with yeast, where it enabled a higher dynamic range and lower leak in trans-kingdom signaling. We hypothesize this is due to leaky translation through the theophylline inducible riboswitch in *S. elongatus*, which might be improved by either more tightly controlled riboswitch elements or via tuning the basal expression of the riboswitch-containing mRNA in future studies.

Our results show that the auxin signals generated by *S. elongatus* can be connected to gene expression in yeast in co-culture via auxin induced degradation. The observation that auxin signals generated similar responses in yeast in co-culture compared to the addition of supernatant implies that the proximity of partners does not impact the efficiency of trans-kingdom signaling. However, future studies that more deeply interrogate the impact of spatial separation are needed. The ease of coupling auxin signals to gene expression via degron fusion enables flexible connection of auxin signals produced by *S. elongatus* to yeast behaviors. Prior work has also identified degron variants that enable a range of different auxin sensitivities and response dynamics, enabling tunable transduction properties^8^. Thus, this system will facilitate future fundamental studies of trans-kingdom microbial interactions. Our group aims to leverage this system for the development of synthetic lichen in the future^1^. This system also has significant translational potential because of its capacity to enable coordinated bioproduction in microbial consortia. Prior studies have shown that *S. elongatus* can be engineered to maintain the growth of co-cultured heterotrophs like yeast, which are engineered for bioproduction, by donating a portion of its fixed carbon as sucrose^16,17^. The auxin-based signaling system we demonstrate here could be applied to enable coordinated behaviors in co-culture, such as growth or bioproduction, potentially improving performance.

The demonstration that auxin signals generated by *S. elongatus* associated with plant roots can modulate the regulation of synthetic transcription factor-based genetic circuits in the plant shows that this trans-kingdom signaling system can be generalized to other species pairs. However, this highlights a limitation of this system as well. In receiver organisms that biosynthesize auxin themselves, such as plants and some fungi, there is a possibility of confounding cross talk. One strategy around this is to tune the biosynthesis in the sender and the sensitivity of the receiver to be outside of physiological ranges, as was done in our plant experiments. However, auxin-based trans-kingdom signaling systems are best applied to organism pairs deficient in auxin signaling to enable utilization of the entire dynamic range of the signaling system. Our experiments involving root-associated *S. elongatus* also serve as proof-of-concept for how microbes could be engineered to enable photosynthesis-driven auxin application to plants. Prior studies have shown that auxin producing root-associated microbes can enhance growth, highlighting the future utility this approach might have for sustainable agriculture^20^.

Thus, our work lays the foundation for the development of synthetic auxin-based trans-kingdom signaling systems that are tunable and can be broadly applied. It demonstrates that such systems are possible across multiple species pairs and identifies constraints on their function which could be improved in future iterations. In this way this work enables future fundamental studies of intraorganismal interactions as well as engineering of multi-kingdom consortia for bioproduction and agriculture.

## Methods

### *S. elongatus* strain construction

Gibson assembly was used to assemble the expression cassettes for either *iaaH* alone or both *iaaH* and *iaaM* driven by a theophylline inducible promoter^13^ into a previously validated shuttle vector that contained an expression cassette for kanamycin resistance as well as homology regions for genomic integration into the neutral site 2 safe harbor locus^21^. The sequences of the plasmids generated can be found in **Supplementary Table 1.**

### *S. elongatus* growth conditions for metabolic characterization

*S. elongatus* cultures were grown in BG11 medium supplemented with 1 g L^−1^ HEPES to a final pH of 8.3 with NaOH. Flasks were cultured in a Multitron incubator (Infors HT) at 32 °C under ambient air CO_2_ or supplemented with 2 % CO_2_ with ∼150 μmol photons m^−2^ s^−1^ of light provided by Sylvania 15 W Gro-Lux fluorescent bulbs and shaken at 150 rpm. Cultures were back-diluted daily to an OD750 of 0.3 and acclimated to the medium/irradiance for at least 3 days prior to experiments Kanamycin (Kn; 12.5 μg mL−1) was used to maintain the stains. For theophylline concentrations from 0.005 to 1 mM. In all cases, antibiotic selection was removed prior to conducting the reported experiments to minimize unintended effects.

### Chlorophyll a extraction

We adapted the established protocol from Lichtenthaler et al.^22^ to measure chlorophyll content. Briefly chl a was extracted from 1 mL of cyanobacteria pellet centrifuged at 16000×g for 10 min and then resuspended in 1 mL 100 % MeOH. Cells were extracted for 30 min at 4°C and vortexed at high-speed vortex for 2 min. Cells were pelleted and absorbance was taken at 665 nM to estimate chl a as previously published^23^.

### Auxin extraction and LC–MS analysis

Cyanobacterial cultures were typically seeded at OD750 = 0.3 with the indicated theophylline concentrations (from 0.005 mM up to 1 mM) with indole-3-acetamide and indole (100 μM each) were used as inducing compounds. Cells were pelleted by centrifugation of 0.5 mL samples at 13,300×g for 10 min, and the supernatant was kept for the extraction. Where appropriate dilution of the supernatant was performed with BG11 before the extractions. To quantify the auxin (IAA) in the supernatant, 0.5 mL of ethyl acetate supplemented with 100nM of C13 IAA as internal standard. 0.5 mL of the extraction solvent and 0.5 mL supernatant was vortex-mixed for 3 min, centrifuged for 1 min and the organic phase (top) was collected. The extraction procedure was repeated 2 times with just ethyl acetate, and the volume extracts (1.5 mL total) were dried using a Speedvac vacuum concentrator (Thermo Scientific). The samples were reconstituted in 500 μL of 50% methanol/water, and 100 μL was immediately transferred to LC–MS vials (with glass inlets) and stored at −20 °C prior to LC–MS/MS analysis, resulting in a final internal standard concentration of 100 nM.

Samples were analyzed on a Waters Xevo TQ-XS tandem quadrupole LC-MS/MS system. 5 uL of sample was injected onto a Waters Acquity UPLC BEH-C18 column (2.1×50 mm) and compounds were separated using the following gradient: initial conditions were 98% mobile phase A (water + 0.1% formic acid) and 2% mobile phase B (acetonitrile) and held for 0.5 min followed by a ramp to 70% B at 3 min, then to 99% B at 4 min, hold at 99% B until 5 min then return to 98% A at 5.01 min and hold until 6 min. Column temperature was 40°C and the flow rate was 0.4 ml/min. Compounds were ionized by electrospray ionization operating in positive ion mode with a capillary voltage of 1.0 kV, source temp of 150°C, desolvation temp of 400°C, cone gas flow of 150 L/hr and desolvation gas flow of 800 L/hr. MS/MS parameters are listed in the table below. Data were processed using the targetlynx tool in Waters Masslynx software. For quantification purposes, standard curve of IAA was quantified and created in the range of 0.96–2000 nM using the same internal standard as mentioned above. Each point in the calibration curve consisted of the same internal standard concentration 100nM C13-IAA as the samples. The ratios of LC– MS/MS peak areas of the analyte/internal standard were calculated and used to construct calibration curves of the peak area ratio against the analyte concentration using unweighted linear regression analysis. For the calibration curve standard IAA was commercially available.

**Table:** MS/MS parameters of each compound.

| compound | Parent ion | Daughter ion | Dwell time (sec) | Cone voltage | Collision energy |
|---|---|---|---|---|---|
| IAM | 175 | 130 | 0.063 | 30 | 16 |
| IAA | 176.1 | 130.1 | 0.063 | 30 | 16 |
| 13C6-IAA (IS) | 182.1 | 136.1 | 0.063 | 30 | 20 |

### Growth Conditions for Co-culture

*S. elongatus* strains with theophylline inducible *iaaH* expression were grown in BG11 medium supplemented with 1 g L^−1^ HEPES to a final pH of 8.3 with NaOH, in a chamber with 50 μmol/m²/s 24-hour light period shaking at 125 rpms maintained at 30°C. Flasks contained a sterile foam plug with a bubbler unit at a flow rate of 1L/-min per L of media supplemented with >1% CO_2_ (CO2Meter). *S. elongatus* cultures were grown for up to a week in advance, grown in 17.5 ng/mL of kanamycin. At the beginning of the co-culture experiment, *S. elongatus* were diluted to 0.3 OD750 in BG11 without antibiotics with freshly prepared theophylline, IAM, or vehicle(water or DMSO), then incubated in the conditions described for 72 hours. 16 hours before the experiment, a single colony of the *S. cerevisiae* containing the Venus-IAA17^19^ was inoculated into 4 mL of synthetic complete media (SCM)^19^ and shaken at 250 rpm at 30°C. At 72 hours, 10 mLs of the *S. elongatus* was collected in a tube, and a 50 mL conical was spun down for 5 minutes at 4000 rpm at room temperature, and the supernatant filtered through a 0.22 µM pore. The *S. elongatus* and the *S. cerevisiae* cell concentration were determined by the flow cytometer and combined at a ratio of 8×10^6^ to 2×10^6^ total cells. The volume was made up to 4 mLs by either the induced or vehicle filtered supernatant. Indole-3-Acetic Acid (IAA) was added during the time of mixing as separate, positive control with vehicle supernatant as the diluent. The mixed *S. elongatus* and *S. cerevisiae* were returned to yeast growth conditions for 3 hours before reading out on the flow cytometer.

### Flow Cytometry

All flow cytometry measurements were acquired using a BD Accuri6™ Plus with attached CSampler using the 488 nm excitation lasers (FL-1:YFP) and 510/15 and (FL3:RFP) 610/20 emission filter Version 1.034.1 Build 20180111.34.1. 10,000 events of mixed cultures were collected using a freshly drawn gate around the *S. cerevisiae* population and exported as FCS files for analysis. Briefly, FCS files were loaded using FCMeasurement class from FLowCytometryTools then populations were plotted using forward and side scatter. Matplotlib’s PolygonSelector and Path(^24^) classes were used to create a custom gating scheme for drawn around uninduced vehicle control and applied to every sample in a biological experiment. Mean fluorescence intensity was collected then normalized by dividing the mean of YFPRFP value for each event. Biological replicates were acquired on different days. Data is accessible on the **Supplementary Files** and a single, representative experiment is plotted.

### Arabidopsis thaliana luminescence assays and confocal microscopy

*A. thaliana* seeds expressing the auxin inducible synthetic transcription factor were surface sterilized and selected on 0.5x MS plates with kanamycin using a previously published light shock protocol^11^. In tandem both wildtype *S. elongatus* as well as the strain with theophylline inducible *iaaH* expression were cultured for 72 hours with 100 µM IAM and 0.5 mM theophylline. A week post-germination the plants were transferred to fresh plates, and their roots were inoculated with 100 µl of *S. elongatus* culture. These were allowed to associate over 24 hours of vertical growth in long day conditions. Following this one set were treated with a solution of 5µM D-luciferin from GoldBio in sterile water, incubated in the dark for 15 minutes, and then the bioluminescence signal was measured with a 1 minute long exposure using an AnalyticJena whole organism imager. ImageJ^25^ was used to quantify the bioluminescence signal in root tips and the elongation zone from these images. The same experimental set up described above was also used to obtain confocal microscopy images. Here 24 hours after treatment with the *S. elongatus* lateral root tips were excised and mounted onto slides and then imaged using a 63x water dipping objective on a Leica confocal microscope, such that the stack went from the middle of the root to its surface. These 3D volumes were then analyzed using the freterantor2^26^ plugin for FIJI, which enabled segmentation of the individual nuclei and measurement of the average fluorescence signal.

### Data analysis and visualization

All collected data was analyzed and plotted in ipython notebooks using jupyterlab. The seaborn package was used to make plots^27^. Additionally, T tests in figures 5 were performed using the statistics package in python. In Figure 4 for ease of visualization, the flow cytometry data is plotted as a smoothed histogram of the cell counts at different fluorescence levels using the kdeplot function in seaborn. This creates a kernel density plot by estimating the continuous probability density function of the dataset, where the total area under the curve equals 1. All code for data analysis and visualization reported in this work, along with the raw data, was compiled into a jupyter notebook in the following github repository (https://github.com/arjunkhakhar/Auxin-based-transkingdom-signaling)

## Abbreviations

IAA: Indole-3-acetic acid, auxin
IAM: Indole-3-acetamide
AFB2: Protein AUXIN SIGNALING F-Box 2
IAA17: *INDOLE ACETIC ACID-INDUCED PROTEIN 17*

## Author Information

Research was performed at: Colorado State University Biology Building 251 W. Pitkin Street Fort Collins, CO 80521 United States of America

Michigan State University Plant Biology Building 612 Wilson Road East Lansing, MI 48824 United States of America

## Author contributions

M.F. designed research, performed research, analyzed data, and revised the paper; E.J.K. designed research and performed research; D.K. designed constructs; B.T.L. performed research; D.C.D. acquired funding, designed research, and revised paper; A.K. acquired funding, designed research, performed research, and wrote paper.

## Acknowledgements

We would like to thank A.K. and D.C.C for their time and reagents which were funded by National Science Foundation grants 2334680 and grant 2334681.

