## Supplemental Information for "Developing synthetic auxin-based trans-kingdom signaling"

| Component | Link to Benchling |
|---|---|
| *Theophylline riboswitch,* iAAH | P360 - MSU_NS2_iaaH · Benchling |
| *Theophylline riboswitch*, **iaaM,** iAAH | *P1043 - MSU_NS2_pTheo_iaaM_pTheo_iaaH · Benchling* |

**Supplementary Table 1**

Table providing the engineered plasmids containing the *iaaH* and *iaaM* genes that were transformed into *S.elongatus*.

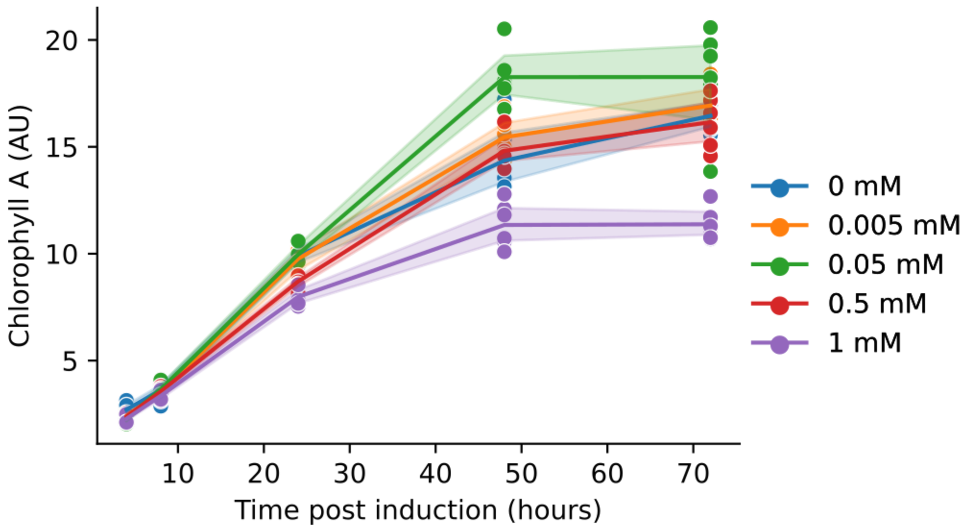


**Supp Fig 1**

*S. elongatus* strain containing theophylline induced expression of *iaaH.* Plot summarizing chlorophyll a (chla) as a proxy of photosynthetic capacity over time, measured as arbitrary units, in media containing 100 µM IAM post induction with varying concentrations of theophylline. Shaded area around lines connecting means represent the 95% confidence interval (n=6).

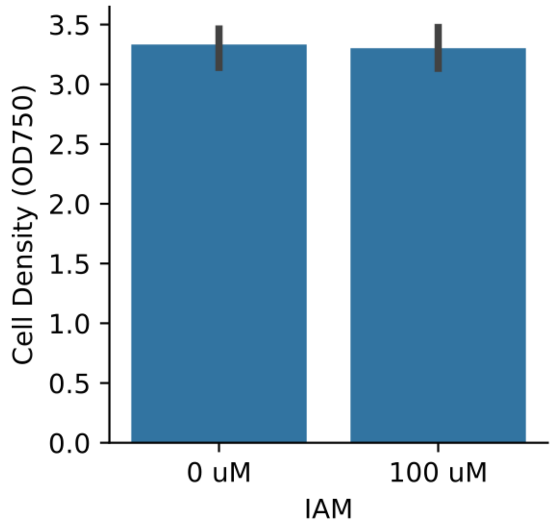


**Supp Fig 2**

*S. elongatus* strain containing theophylline induced expression of *iaaH.* Bar plot summarizing cell density at 48 hours, measured as OD750, in media containing 100 µM IAM and no theophylline (n=1).

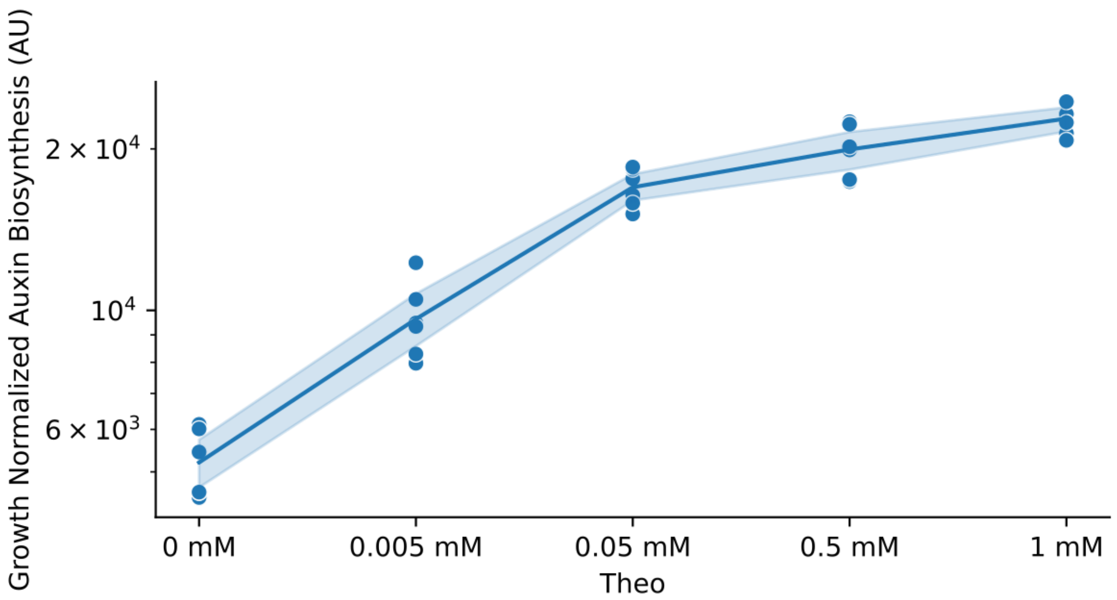


**Supp Fig 3**

*S. elongatus* strain containing theophylline induced expression of *iaaH.* Plot summarizing auxin normalized by cell growth measured as arbitrary units (AU). Cell density measured as OD750, and auxin measured by LC-MS in media containing 100 µM IAM 48 hours post induction with varying concentrations of theophylline. Shaded area around lines connecting means represent the 95% confidence interval (n=6).

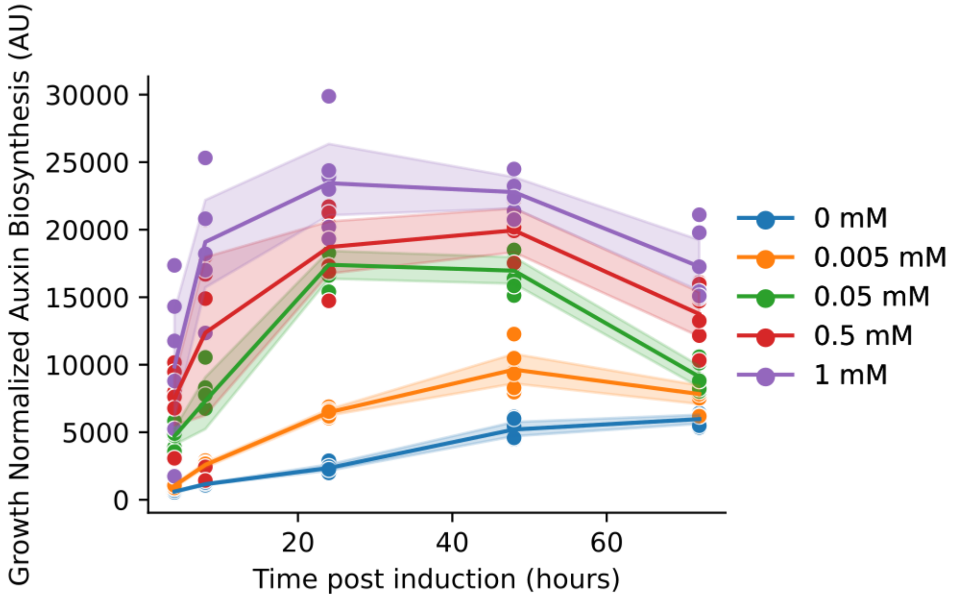


**Supp Fig 4**

*S. elongatus* strain containing theophylline induced expression of *iaaH.* Plot summarizing auxin normalized by cell growth over time, measured as arbitrary units (AU). Cell density measured as OD750, and auxin measured by LC-MS in media containing 100 µM IAM 48 hours post induction with varying concentrations of theophylline. Shaded area around lines connecting means represent the 95% confidence interval (n=6).

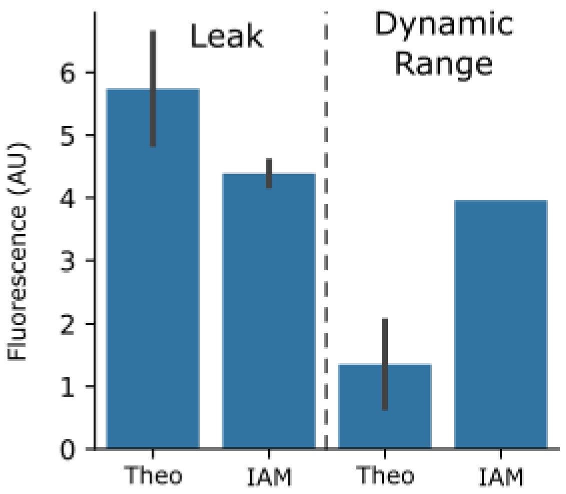


**Supp Fig 5**

Bar plots summarizing mean normalized YFP fluorescence of approximately 10,000 yeast cells per independent replicate treated with supernatant from cultures of *S. elongatus* with inducible auxin biosynthesis that were incubated with IAM, DMSO, or theophylline. Leak (left) represents vehicle treatment minus either 1 mM theophylline (Theo) or minus 100 uM IAM (IAM). Dynamic range (right) represents supernatant with a fixed concentration of IAM (100uM) with no theophylline minus IAM with 1 mM theophylline (Theo) and theophylline (1 mM) with 100 uM IAM minus theophylline with 500 uM IAM (IAM). The errors bars represent 95% confidence interval of n =2.
